# Duesselpore™: a full-stack local web server for rapid and simple analysis of Oxford Nanopore Sequencing data

**DOI:** 10.1101/2021.11.15.468670

**Authors:** Christian Vogeley, Thach Nguyen, Selina Woeste, Jean Krutmann, Thomas Haarmann-Stemmann, Andrea Rossi

## Abstract

Genome-wide analysis of transcriptomes offers extensive insights into the molecular mechanisms underlying the physiology of all known species and discover those that are still hidden. Oxford Nanopore Technologies (ONT) has recently been developed as a fast, miniaturized, portable and a cost effective alternative to Next Generation Sequencing. However, RNA-seq data analysis software that exploit ONT portability and allows scientists to easily analyze ONT data everywhere without bioinformatic expertise is not widely available. We developed Duesselpore™, an easy-to-follow deep sequencing workflow that runs as a local webserver and allows the analysis of ONT data everywhere without requiring additional bioinformatic tools or internet connection. Duesselpore™ output includes differentially expressed genes and further downstream analyses, such as variance heatmap, disease and gene ontology plots, gene concept network plots and exports customized pathways for different cellular processes. We validated Duesselpore™ by analyzing the transcriptomic changes induced by PCB126, a dioxin-like PCB and a potent aryl hydrocarbon receptor (AhR) agonist in human HaCaT keratinocytes, a well characterized model system. Duesselpore™ was specifically developed to analyze ONT data but we also implemented NGS data analysis. Duesselpore™ is compatible with Microsoft and Mac operating systems, allows convenient, reliable and cost-effective analysis of ONT and NGS data.

## Introduction

In the past decade, RNA-Sequencing (RNA-Seq) became the leading method to analyse whole genome transcriptomes and was more and more implemented into modern medicine (1). At this, RNA-Seq is used in diagnosis, prognosis and therapeutic selection in the fields of infectious disease, fetal monitoring and cancer (1, 2). Additionally, the ability to sequence DNA and RNA quickly and inexpensively is essential to scientific research (3). This necessity has prompted the development of many sequencing techniques beyond the original Sanger sequencing (4). The rise of next-generation sequencing (NGS) has considerably improved the output of data generated by Sanger sequencing (first-generation sequencing) (5). NGS approaches have become widespread in research and diagnostic laboratories and have greatly increased our knowledge of many genetic disorders (4, 6–9). Nevertheless, these technologies remain expensive, laborious, time-consuming, and affected by the limits of short-read sequences generation. As some kind of a new generation, new sequencing methods were established, which aim to sequence long DNA or RNA molecules (10) including Oxford Nanopore Technologies (ONT).

ONT sequencing is based on transmembraneous proteins (nanopores) embedded into a lipid membrane and measures changes in electric current across these pores. These changes are caused by nanosized molecules, such as DNA or RNA (or even aminoacids), as those occupy a volume that interferes with the ion flow and can be recorded by a semiconductor-based electronic detection system and afterwards translated into sequences (11, 12). In the last few years, ONT sequencing has proven to be a fast and cost effective alternative towards other sequencing techniques. Even if the read quality does not reach the high standard of other techniques, it offers other advantages such as fast and easy library preparation, real-time sequencing, PCR-free, direct RNA and ultra long read sequencing (13, 14). (13, 14). The latter is especially useful to improve de novo assembly, mapping certainty, transcript isoform identification, and detection of structural variants (15). Additionally, some ONT sequencing devices are very small granting portability and making sequencing experiments possible outside the laboratory. Nevertheless, sequencing analysis produce large raw datasets that require bioinformatic skills, familiarity with statistics and a Linux-based system to decode these informations. Available software to analyse those data lack a clear and friendly user interface and are mainly limited to basic analysis.

Besides, each RNA-Seq workflow runs on a single method which may bias the result. Therefore, reproducing other experiments needs many installation and validation steps to make it compatible with other pipelines. Moreover, the RNA data processing workflow uses dozens of dependent libraries and third party software. Each of this required package has to follow a strict version of compatible matrices and is designed to run on a Linux operating system. Additionally, most users are not willing to upload their unpublished data to an open platform to employ open-access web services. A client-based computational approach implemented in Javascript is an alternative solution that shifts the computation workload to the client side. Schmid-Burgk and co-workers deployed a lightweight solution to analyze genome editing experiments (16). However, RNA-Seq workflows are more complicated and have not been translated or compiled into lightweight Javascript-like softwares. Moreover, ONT offers mobile platforms that can work in environments, where internet access is limited or absent.

To overcome these issues and to help the community process larger ONT sequencing datasets, we developed a local and mobile web server which we named Duesselpore™, that supports the mobility of Nanopore sequencers. Instead of using a centralized system, we deploy an efficient local-based system that runs on Docker container or a virtual machine (VM) without the necessity to be connected to the internet. Running on a virtual environment has further advantages, as the pipeline runs in an isolated virtual environment and avoids the software to collide with the host’s software.

## METHOD AND SYSTEM DESCRIPTION

### HaCaT cell culture and stimulation

HaCaT keratinocytes were cultivated at 37 °C and 5 % CO_2_ in DMEM medium (PAN Biotech, Aidenbach, Germany) low glucose (1 g/l) supplemented with 10 % FBS and 1 % antibiotics/antimycotics (PAN Biotech). The next day HaCaT keratinocytes were stimulated in full growth medium for 24 h with 1 μM PCB126 or were solvent treated, using 0.1 % DMSO.

### Library preparation

Total RNA was isolated by using the GenUP Total RNA Kit (Biotechrabbit, Hennigsdorf, Germany). The High Sensitivity RNA Screen Tape System (Agilent Technologies, Santa Clara, CA) was used to assess the quality of isolated RNA. 50 ng of total RNA was reverse transcribed and samples were barcoded with the PCR cDNA Barcoding Kit (SQK-PCB109, Oxford Nanopore Technologies, Oxford, United Kingdom). Quantity of amplified cDNA was then determined with the Qubit™ 4 Fluorometer (Invitrogen, Carlsbad, CA), and the range of fragment size was examined using the Agilent D1000 SreenTape assay (Agilent Technologies). The FlowCell Priming Kit (EXP FLP002, Oxford Nanopore Technologies) was used to prime the Flowcell (FLO-MIN106), and an equal amount of barcoded cDNA was loaded. Sequencing was carried out with a MinION (MN33710) using the MinKNOW software (v.21.02.1) over a period of 72 h.

### Basecalling

Raw fast5 reads were base-called and demultiplexed using Guppy (v4.5.4+66c1a7753) using the following command. By this, reads with a quality score below 7 were excluded.

~~~
guppy_basecaller --min_qscore 7 --trim_barcodes --barcode_kits “SQK-PCB109” --compress_fastq -i {input.fast5} -s {output.folder} -cdna_r9.4.1_450bps_hac.cfg --x auto --chunks_per_runner 128
~~~

Afterward, the resulting fastq files of each sample were concatenated into one file.

### Input data and run parameters

Our web service validates the input first by automatic format detection. Input fastq files are compressed and structured in each subdirectory named by experimental condition. Each subdirectory contains multiple replicas of each condition. All other parameters are set as optimal for typical long read RNAseq.

### Quality control

Our RNASeq workflow follows the standard data processing pipelines. First, we employ the quality control with FastQC to quantify the sequence quality and export the quality into HTML files.

### Align and assign read to gene/transcriptome

We use the cutting edge tool Minimap2 (17) to map our read into reference genome/transcriptome. Aligned BAM files are processed by Rsubread/featureCounts (18), HTSeq/htseq-count (19), or Salmon (20) to generate the raw count read gene or transcriptome. Although Salmon can quantify directly from fastq to reference genome, Minimap2 shows a higher assigned ratio for handling noisy ONT data.

### Differential expression analysis and gene ontology

The raw read counts matrices are processed by the difference expression method DESeq2 (21), edgeR/limma (22, 23). DESeq2 shows the high capability to process data with a small number of replicas, which are common issues in NGS.

To analyze Gene Ontology, we integrate the Gene Ontology pipeline using Bioconductor. We cluster our results into various gene ontology pathways such as Gene Set Enrichment Analysis of Gene Ontology (gseGO), Disease Ontology, Network of Cancer Gene (DOSE) (24), Pathview (25).

### Web Server and backend platform

Our web server is implemented in Python 3.8 with Django 3.2 web framework which has long-term support from the open-source community. We use bio-conda, Biopython, and R/Bioconductor run on the Ubuntu 18.04 system packed in Docker container or Virtual machine (VM) as backend. We implement swap memory in VM version to extent our platform memory which is required for running high memory consumption software such as minimap on an average configuration PC. The Docker or VM is elastic, which can be configured depending on the configure of the host PC. All software is free and open-source. The detailed system architecture and manual are described in the Supplementary document.

## Results

### Validation of Duesselpore™

To validate Duesselpore™, immortalized HaCaT keratinocytes were stimulated with 3,3’,4,4’,5-pentachlorobiphenyl (PCB126) or were solvent treated with 0.1 % DMSO for 24 h. PCB126 was chosen as the mode of action, because how it regulates gene expression is well understood. According to their lipophilic properties, PCB126 molecules can easily pass through the cell membrane and bind the aryl hydrocarbon receptor (AHR), a cytosolic transcription factor trapped in a multiprotein complex. Upon ligand binding, the multiprotein complex dissociates and the AHR translocates into the nucleus, where it dimerizes with the aryl hydrocarbon receptor nuclear translocator (ARNT) and binds to xenobiotic responsive elements (XRE) in the genome This signalling pathway induces the expression of various genes encoding for Phase I and Phase II biotransformation enzymes (26). Amongst these genes are the members of the superfamily cytochrome P450 (CYP) *CYP1A1, CYP1B1* and aldehyde dehydrogenases (ALDH) *ALDH3A1* as well as for proteins involved in cell proliferation, differentiation and apoptosis (27).

Total RNA was isolated and transcribed into complementary DNA (cDNA) as described. cDNAs were barcoded, loaded in the same flowcell and sequenced over a period of 72 h to ensure a sufficient number of reads for further downstream analyses. The raw .fast5 output files were basecalled with *Guppy*, ONTs basecalling algorithmand and demultiplexed (barcoded sorted). As it requires a tremendous amount of computational resources, especially parallel computation run on the graphic card, this step was not included in Duesselpore™ and was run beforehand. Nevertheless, it is worth mentioning that basecalling is offered in accurate or fast mode in MinKNOW, the software used to run an Oxford Nanopore experiment. The generated fastq files were then concatenated and analyzed with Duesselpore™.

Duesselpore™ provides a mapping summary, including the number of reads, how many reads were mapped, and the percentage of mapped reads (Table 1). Secondary mappings were excluded as those are not considered when these sequences are assigned to genetic features.

**Table1:** Mapping summary of sequences aligned with minimap2.

|  | PCB_replicate1 | PCB_replicate2 | PCB_replicate3 | DMSO_replicate1 | DMSO_replicate2 | DMSO_replicate3 |
| --- | --- | --- | --- | --- | --- | --- |
| *nreads* | 4929644 | 1561372 | 3624589 | 2877111 | 2122355 | 4027410 |
| *read mappings* | 4995752 | 1586505 | 3651019 | 2918088 | 2150179 | 4055402 |
| *Secondary* | 0 | 0 | 0 | 0 | 0 | 0 |
| *Supplementary* | 66108 | 25133 | 26430 | 40977 | 27824 | 27992 |
| *Duplicates* | 0 | 0 | 0 | 0 | 0 | 0 |
| *%mapping* | 87.66 | 92.94 | 75.96 | 92.93 | 89.88 | 78.79 |

A more detailed samples overview is provided in the sample summary table, which contains information about the read length, million base sequences (mbs), mean quality score (qval), and GC content (gc) (Figure 2A). Additionally, read length and quality score distributions are depicted in violin plots (Figure 2B and C).

**Figure1:**
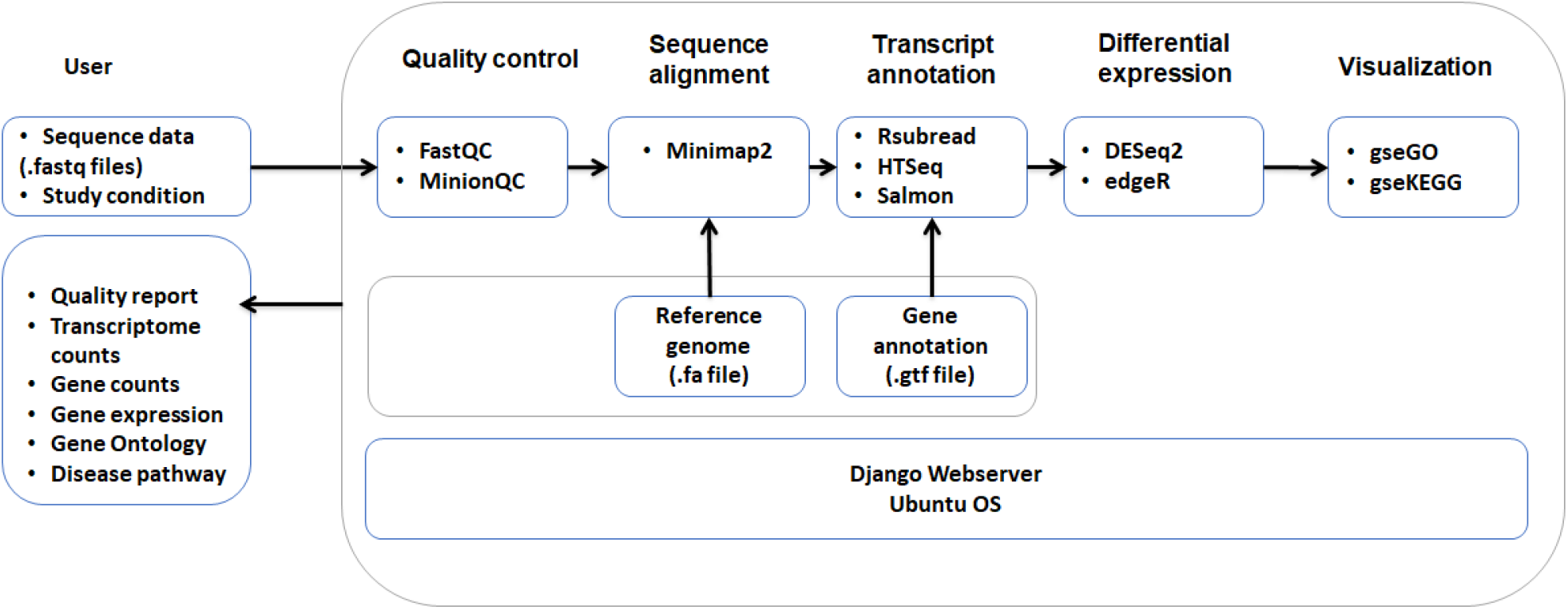
Workflow and system description

**Figure 2:**
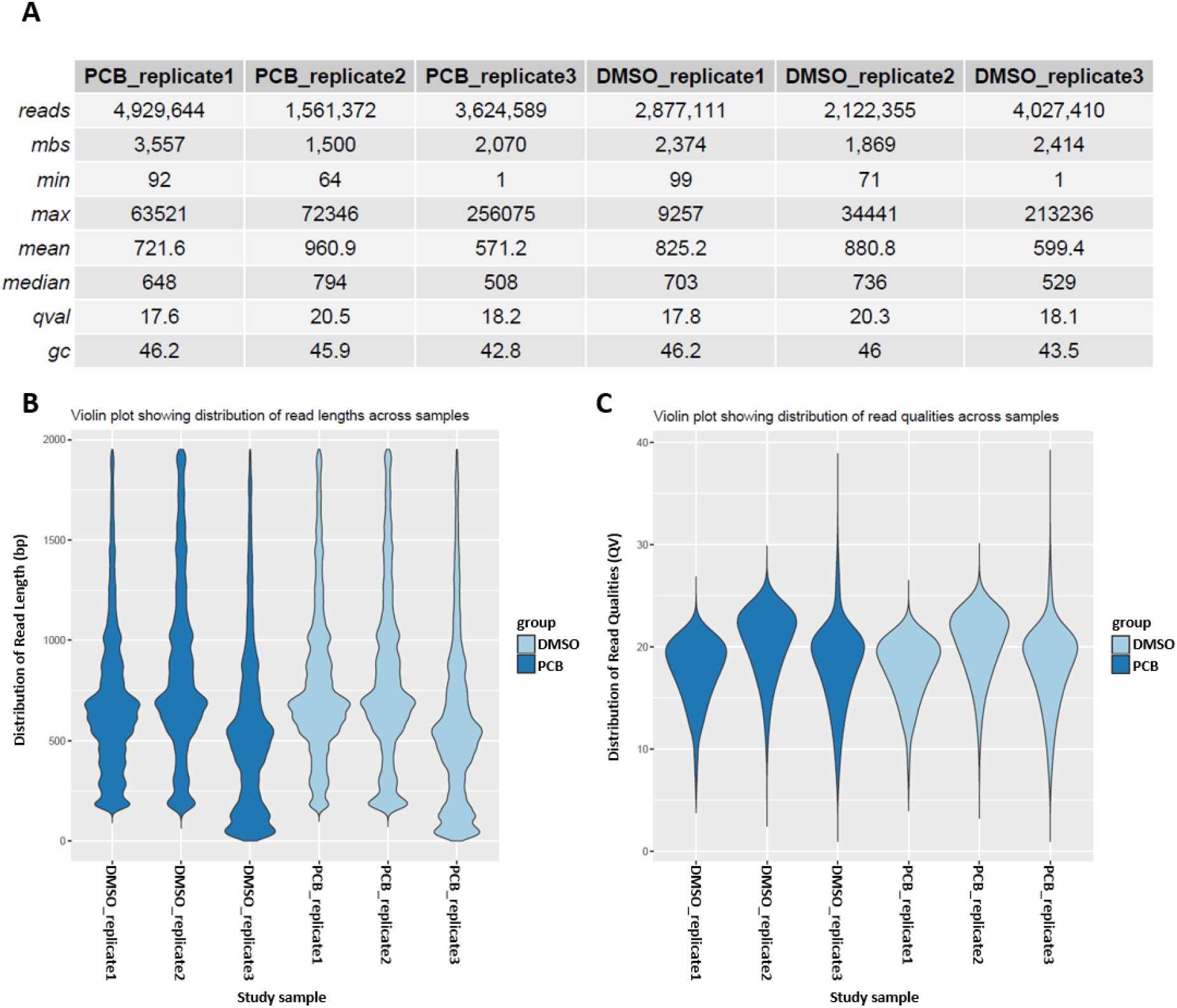
Sample summary and quality control. (A) Summary of the reads of every sample. The table gives information about the number of reads, million base sequences (**mbs**), minimal and maximum read length, mean and median read length, quality score (**qval**) and GC content (**gc**). Violin plots depict the distribution of read length (B) and the distribution of read qualities (C) across different samples.

After carrying out sample quality control, Duesselpore™ performs differential gene expression (DGE) analysis. Thereby, the user is able to choose between the packages *DESeq2* or *edgeR*. Both packages use statistical models based on the negative binomial distribution and are the most commonly used tools. It is recommended to use *DESeq2* when the number of replicates is relatively low (below 5) as this tool shows the highest consistency of the identification of significantly differentially expressed (SDE) genes and shows the lowest false discovery rate (28). The results of the differentially expressed gene (DEG) analysis are provided as a excel file, and in our data set, the genes *CYP1A1, CYP1B1, ALDH3A1* as well as other established AHR target genes, such as ATP-binding cassette super-family G member 2 (ABCG2), plasminogen activator inhibitor-2 (SerpinB2) and TCDD-inducible poly [ADP-ribose] polymerase (TiPARP), are among the SDE genes. The regulation of *CYP1A1, CYP1B1*, and *ALDH3A1* was confirmed by qRT-PCR (data not shown). The top 30 genes with the highest variance are depicted in a variance heatmap (Fig. 3A), also provided by Duesselpore™. For this purpose, the counts per gene were normalized to the counts per million (CPM) scaling factor (23). The number of depicted genes can be defined in the user interface.

**Figure 3:**
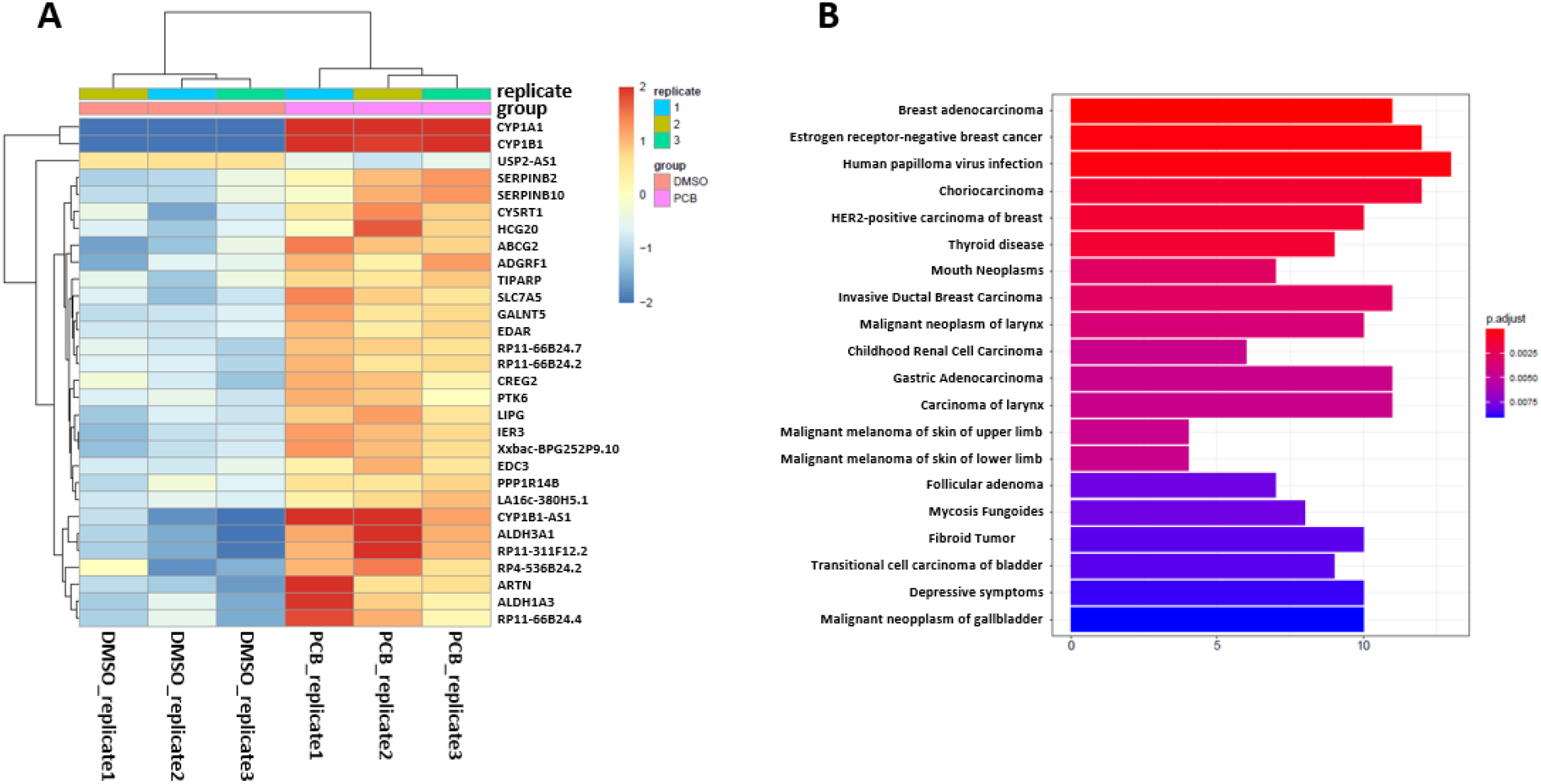
Variance of gene expression and enrichment analysis. (A) Heatmap showing the top 30 genes with the highest variance after normalizing for the CPM. (B) Enrichment analysis based on the DisGeNET database, revealing diseases, which are associated with the differentially expressed genes. X-axis shows the number of genes involved in each disease.

Besides the detection of differentially expressed genes, Duesselpore™ will also conduct gene set enrichment analyses (GSEA), enrichment analysis based on the DisGeNET (29) and pathway-based data integration and visualization focusing on KEGG (Kyoto Encyclopedia of Genes and Genomes) pathways (25). These features will help the user to put the data into context and to identify potential signaling pathways and diseases based on the differentially expressed genes. To select a certain KEGG pathway, the user needs to provide a KEGG identifier and enter it into the webform. PCB126 is highly toxic and exhibits mutagenic potential, which is not solely based on its capability to activate the AHR (30). Hence, genes regulated as a consequence of PCB126 exposure were associated with various malign neoplasias according to the DisGeNET database (Fig. 3B). The relation between genes and disease is depicted in a network plot, either in a non-circular (SFig. 1A) or circular (SFig. 1B) format.

Duesselpore™ outputs were successfully validated by using further datasets (data not shown). Here, the ability to detect PCB126 induced gene expression proves the functionality of Duesselpore™.

## Discussion

Over the last decade, RNA-sequencing methods became more affordable and advanced to a commonly used technique to analyze genome-wide transcriptomes. While each step of sequencing has been drastically simplified, i.e. sample preparation due to the implementation of preparation kits, users are still overencumbered by the size and the amount of raw data, that require further analysis. These analysis often need bioinformatic expertise and a certain amount of computational resources. In order to reduce this burden, we developed Duesselpore™, a local web server that delivers a solution without interfering with the mobile characteristics of ONT sequencers, such as no internet connection. To ensure its functionality, Nanopore reads were generated using DMSO or PCB126 treated HaCaT keratinocyte samples and analyzed using Duesselpore™. Among the DEGs, many prototypic genes that are regulated by the AHR, such as *CYP1A1, CYP1B1, ALDH3A1* or *SERPINB2*, were upregulated in the test data set. The gene set enrichment analysis revealed an upregulation of gene patterns, which are connected with different malign neoplasias. This observation is associated with the mutagenic potential of PCB126 and the fact that the AHR activity is enhanced in many different tumour entities (31). These results were expected and thus, validate the data obtained by Dusselpore™ analysis (26, 30).

Duesselpore™ interface allows the selection of five commonly used genomes (human, rat, mouse, zebrafish and c. elegans), provides different pipelines to analyze RNA-sequencing data, and by using publicly available tools, it helps users to perform advanced bioinformatical data analysis, generates figures and tables that are suitable for publication. All these tools require various dependencies, we encapsulated and compiled this workflow into two version: a Docker image and a virtual machine, making Duesselpore™ independent from the host’s machine. All these properties grant the user a strong flexibility in analysing data even without profound bioinformatical knowledge. Furthermore Duesselpore™ is not limited to the analysis of ONT derived data but it also allows the analysis of Illumina and PacBio data (Supplementary Files).

In summary, we developed and successfully validated Duesselpore™ as a webtool, which is able to analyse in detail ONT and NGS (PacBio or Illumina) derived RNA-Seq data without the requirement of a profound bioinformatical background. We expect Duesselpore™ to be of great interest to the scientific community.

## Declarations

### Code availability and data

Please read Duesselpore manual for more information and software links

https://github.com/thachnguyen/duesselpore

Docker image.

https://hub.docker.com/repository/docker/thachdt4/duesselpore

Test dataset (DMSO PCB standard and lightweight):

- https://iufduesseldorf-my.sharepoint.com/:u:/g/personal/thach_nguyen_iuf-duesseldorf_de/EWIk4CLauThHk61_5rItjEcBrSUl1a3_oZ6QxjfdDmdqsA?e=kRPvaa
- https://iufduesseldorf-my.sharepoint.com/:u:/g/personal/thach_nguyen_iuf-duesseldorf_de/ES4BsdfJSKNHl-mDUR3BogcB1HpDzV7eZ1kTikAd2Esq8w?e=lfDF9E

Sample result from ONT

https://github.com/thachnguyen/duesselpore/blob/main/sample_result/nanopore/sample_result_ONT.zip

Sample result from Illumina

https://github.com/thachnguyen/duesselpore/raw/main/sample_result/illumina/test_illumina.zip

Sample result from PacBio

https://github.com/thachnguyen/duesselpore/blob/main/sample_result/pacbio/Test_Pac_bio.zip

### Ethics approval

Not applicable

### Consent to participate

Informed consent was obtained from all individual participants included in the manuscript

### Consent for publication

Consent for publication was obtained from all individual participants included in the manuscript

### Author contributions

AR conceptualized this work; CV and TN programmed the webtool and performed data analysis; SW conducted library preparation and sequencing; THS provided resources; AR, CV and TN wrote the manuscript; JK and THS revised the manuscript.

---

**Figure S1:**
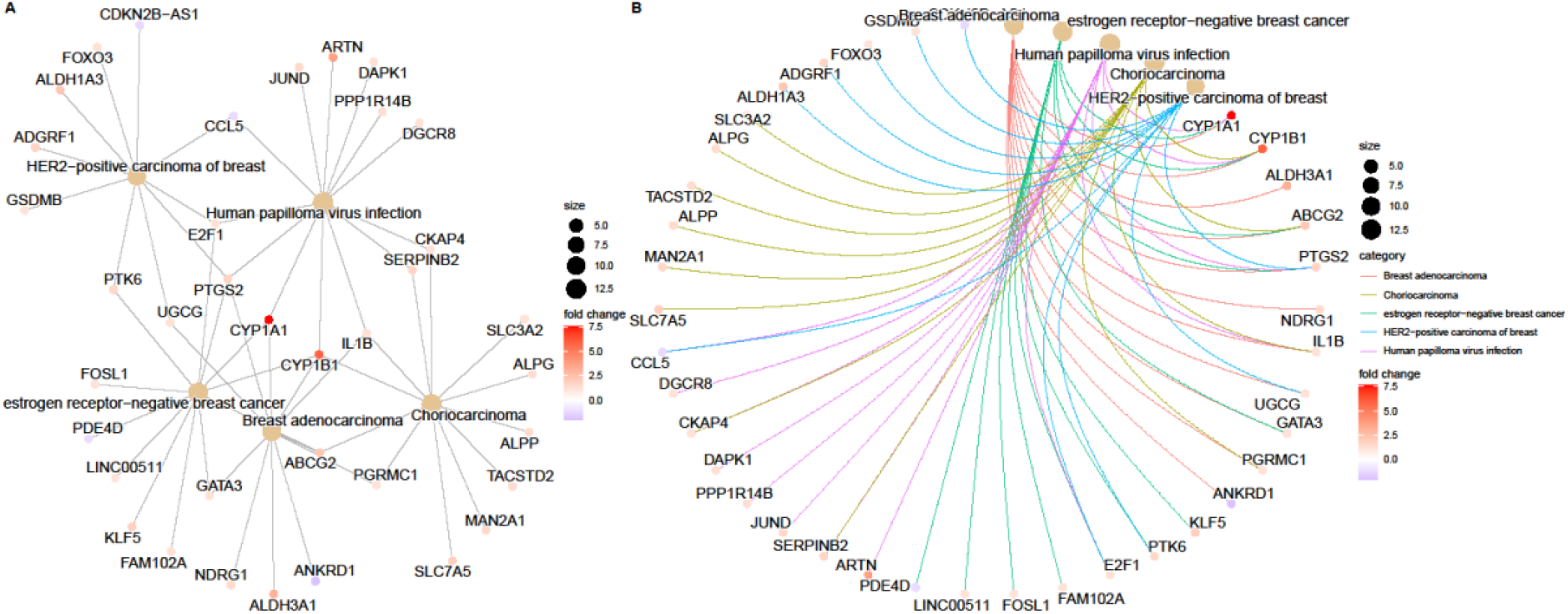
Enrichment analysis according to the DisGENet. Enriched NCG categories are linked with the corresponding genes. Size of the dots depicts the number of genes associated with the respective NCG category and the color code visualizes the log2 fold change of PCB126 treated samples over the DMSO control group. The network is shown in a (A) linear and (B) circular confirmation.

